# Monte Carlo Sampling of Protein Folding by Combining an All-Atom Physics-Based Model with a Native State Bias

**DOI:** 10.1101/361527

**Authors:** Yong Wang, Pengfei Tian, Wouter Boomsma, Kresten Lindorff-Larsen

**Affiliations:** Structural Biology and NMR Laboratory, Linderstrøm-Lang Centre for Protein Science, Department of Biology, University of Copenhagen, Ole Maaløes Vej 5 DK-2200; Copenhagen N, Denmark, Laboratory of Chemical Physics, National Institute of Diabetes and Digestive and Kidney Diseases, National Institutes of Health, Bethesda, Maryland 20892, United States; Department of Computer Science, University of Copenhagen, 2100 Copenhagen Ø, Denmark

## Abstract

Energy landscape theory suggests that native interactions are a major determinant of the folding mechanism of a protein. Thus, structure-based (Gō) models have, aided by coarse-graining techniques, shown great success in capturing the mechanisms of protein folding and conformational changes. In certain cases, however, non-native interactions and atomic details are also essential to describe the protein dynamics, prompting the development of a variety of structure-based models which include non-native interactions, and differentiate between different types of attractive potentials. Here, we describe an all-protein-atom hybrid model, termed ProfasiGo, that integrates an implicit solvent all-atom physics-based model (called Profasi) and a structure-based Gō potential, and its implementation in two software packages (PHAISTOS and ProFASi) that are developed for Monte Carlo sampling of protein molecules. We apply the ProfasiGo model to study the folding free energy landscapes of four topologically similar proteins, one of which can be folded by the simplified potential Profasi, and two that have been folded by explicit solvent, all-atom molecular dynamics simulations with the CHARMM22^∗^ force field. Our results reveal that the hybrid ProfasiGo model is able to capture many of the details present in the physics-based potentials, while retaining the advantages of Gō models for sampling and guiding to the native state. We expect that the model will be widely applicable to study the folding of more complex proteins, or to study conformational dynamics and integration with experimental data.

## Introduction

It is an essential biological fact that most,^1^ though not all,^2^ naturally-occurring proteins can self-organize to ordered three-dimensional structure(s). There has thus been an enormous progress in solving protein structures, as evidenced by the observation that the Protein Data Bank has collected more than 142,000 structures up to date. Despite substantial progress in combining experiments, theory and simulations to study protein folding,^3–9^ there is, however, a substantial gap between the number of structures we know and the proteins for which we know the folding mechanism. In addition to the intellectual challenge involved in understanding the protein folding mechanism, modeling protein folding has potential uses in for example protein design,^10,11^ in molecular drug development,^12^ and in interpreting pathogenicity of genomic sequence variation. ^13^

Recent advances in computer hardware, methods for enhancing sampling and protein force fields have made simulations an irreplaceable tool in the study of protein folding.^14–18^ In principle, a long, equilibrium molecular dynamics (MD) simulation based on an accurate all-atom, physics-based, explicit-solvent model can not only provide spatial and temporal details on the structural ensembles of folded states, but also elucidate the mechanism of folding/unfolding transitions.^19^ While this has been achieved for small fast-folding proteins,^5^ and even a natural protein,^20^ such work generally requires access to extensive sampling using specialized hardware (e.g. Anton^21^). Further, most proteins are not ‘fast-folding’,^22^ and although it is possible also to reach long timescales through using e.g. Markov state models^23^ or enhanced sampling techniques,^24^ it will not be possible to study folding processes for many proteins using routine all-atom MD simulations in the foreseeable future.

As an alternative to the detailed all-atom physics-based models, native-structure-based models (also called Gō models^25^) have been widely applied to investigate the folding and assembly mechanisms of ordered and disordered proteins.^26–28^ These models are applicable to larger sizes, complex topologies and slow kinetics, especially when aided by coarse-graining techniques.^29,30^ The success of these models has been explained by the proposal that such models naturally realise a key feature of naturally-occurring proteins, that is, a minimally frustrated and ideally funnel-shaped energy landscape, ^31^ and indeed analyses of all-atom MD simulations reveal the central role of native contacts.^32^ The principle of minimal frustration in energy landscape theory directly leads to a conclusion that protein topology is a key factor governing the folding mechanism.^32–34^ Currently, there are many software tools available to build and simulate Gō-type models, including SMOG,^35^ AWSEM-MD,^36^ CafeMol,^37^ MMTSB,^38^ CHARMMing,^39^ eSBMTools,^40^ NAMD-Go,^41^ SOP-GPU,^42^ GENESIS,^43^ MonteGrappa,^44^ and SIMONA.^45^ Most of them are based on MD simulation though the last two utilise a Monte Carlo (MC) framework. Also, nearly all previously used Gō-type models have employed a coarse grained representation of the protein.

In most Gō-type models, the native interactions are emphasised by an attractive poten-tial, while the interactions not present in the native folded structures (non-native interactions) are usually simply treated with a repulsive potential. Nevertheless, these non-native interactions may have significant impact on folding process by adding ‘roughness’ to the energy landscape,^46–50^ such as trapping in misfolded or intermediate states and causing aggregation and disease.^49^ Residual non-native interactions, resulting in local violations of the minimal frustration principle,^51^ are considered to be a consequence of the conflicting requirements of foldability and function of a protein sequence.^52^ Opposite to the common view that the non-native interactions contribute primarily to the roughness of landscapes and frustrate the folding process, there are also cases that demonstrate that the non-native interactions facilitate the biological process and play an effective role in protein folding.^53–57^ The potentially important role and related open questions of non-native interactions have driven the development of many enhanced structure-based models^58^ by introduction of additional potentials (e.g. the Debye-Hückel-type potential to approximate electrostatic interactions at low salt concentrations,^59–61^ and the Gaussian potential to model hydrophobic interactions ^48,56,62^) and heterogeneous energetic parameters to distinghuish between different types of contacts.^30,60,63,64^

Inspired by previous hybrid models and multi-scale strategies,^65–71^ we have developed a hybrid physics-based and structure-based model (denoted as ProfasiGo model) within the framework of both PHAISTOS^72^ and ProFASi^73^ simulation packages for Monte Carlo simulation of protein molecules. In our model, the physics-based term is inherited from the implicit solvent force field, denoted as Profasi, which has previously been used extensively to study protein folding, aggregation and protein structure determination.^74–78^ (Note that there is both a simulation software package and an energy function called Profasi; we use the term Profasi for the energy function and ProFASi for the software package.) The physics-based term is transferable and preserves the atomistic representation (including hydrogen atoms) of the protein. We then introduce the structure-based potential (*E*_Go_) as an additional term, thus ‘funnelling’ the underlying energy landscape further, so as to accelerate the folding transitions. In this way the hybrid model is designed to be able to capture more complex energy landscapes. In addition, our software architecture facilitates the investigation of the driving force in protein folding (e.g. electrostatic, hydrophobic interactions and hydrogen bonds) through adjustment of the corresponding potential terms.

In this paper we focus on describing and validating the approach in studies of protein folding. In particular we study four α-helical bundles, the designed proteins α3W and α3D, and the homeodomains EnHD and UVF, that pairwise have similar topologies but differ in folding mechanism. One protein can be folded with the pure Profasi force field, and two with the all-atom CHARMM22^∗^ force field, enabling us to examine the extent to which the hybrid model may capture folding mechanisms in a force field without the structure-based term.

## Models and Methods

### Profasi model

The Profasi model belongs to the class of implicit solvent all-atom models (including all hydrogen atoms) designed for MC simulation^73^ and has been applied in protein folding, aggregation, and determination of protein structures and ensembles with experimental restraints.^74,75,77–80^ In Profasi the flexible degrees of freedom are the Ramachandran (*ϕ* and *ψ*) and side chain (*χ*) torsional angles, whereas bond lengths and angles, and peptide plane *ω* torsional angles, are fixed. The interaction potential is composed of four terms:

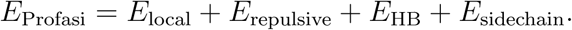

The first term, *E*_local_, accounts for local interactions between atoms, such as the electrostatic interactions between adjacent residues. The other three terms (*E*_repulsive_, *E*_HB_ and *E*_sidechain_) account for non-local interactions: excluded-volume effects, hydrogen-bond interactions, and residue-specific interactions between pairs of side-chains, respectively. The precise form of these four terms can be found in the original description.^75^

### Atomic Gō model

In general, the potential energy function of any Gō-type models, *E*_Go_, as a function of the coordinates of the native structure, Γ_0_, can be simplified into three terms:

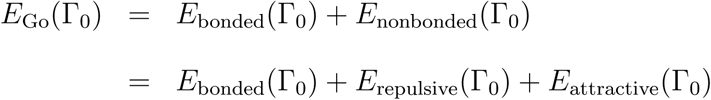

The first two terms, *E*_bonded_ and *E*_repulsive_, maintain the correct backbone geometry, while the last term *E*_attractive_ determines the folding of a peptide chain by attractive inter-atom or coarse grained inter-residue interactions. These attractive interactions are generally defined by a pairwise contact list derived from native structure, called the native contact map. Construction of a native contact map is thus a key step to build a Gō model. In the context of a standard Gō model, the short-range forces to stabilise the native state (e.g. hydrogen bonding, salt bridges and VDW interactions), are approximately represented by the native contact map, while the long-range or nonlocal interactions, like protein-water interactions or water-mediated interactions, are considered to be averaged out and described using a mean field perspective. Currently, there are several algorithms to define the native contact map, including a cutoff-based algorithm,^29,55^ Shadow Contact Map (SCM),^81^ and Contacts of Structural Units.^82^ The native contact maps calculated by different algorithms differ from each other to certain extents, but the resulting thermodynamic properties and folding mechanism are reasonably consistent and robust.^83–85^

For comparison with our hybrid model, we constructed a pure atomic (without hydrogen atoms) Gō model using SMOG^35^ with default parameters (an all-atom contact map by SCM with a cutoff of 6.0 Å). Briefly, SCM is an algorithm to determine contacts between interior protein surfaces without allowing unphysical or occluded contacts.^81^ Here, the allatom attractive Gō potential
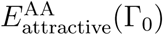 is expressed by a Lennard-Jones (LJ) potential:

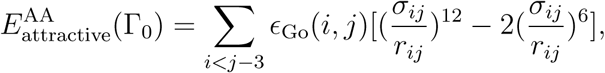

where, *σ_ij_*and *r_ij_* are the native and instantaneous distance between atom *i* and atom *j*, and ϵ_Go_(*i*. *j*) is the strength of pairwise attractive potential between atom *i* and atom *j*. It was homogeneously set to be 1.0 in this work, although it could be tuned to introduce sequence information,^64,86^ through e.g. the Miyazawa-Jernigan matrix^63^ or multi-scaling methods.^87^ Aiming to represent a standard Gō model and for fair comparison, we kept the energetic parameters as general as possible.

### ProfasiGo model

We integrated a structure-based potential (*E*_Go_) into the Profasi force field, and termed this hybrid model ‘ProfasiGo’: *E*_ProfasiGo_ = *E*_Profasi_ + *E*_Go_. We opted to use a coarse-grained version of the native contact map in which only *C_α_*-*C_α_* contacts are included so as to introduce minimal extra potential into Profasi force field. In other words, the Gō potential was introduced as a minimal perturbation. We implemented two variants of *E*_Go_ into both the ProFASi and PHAISTOS software packages (which already implemented the Profasi energy function). One is based on a 12-10 LJ-like potential:

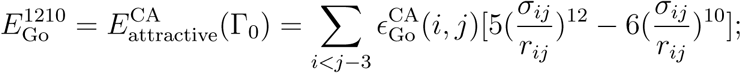

and the other uses a 12-10-6 potential described by

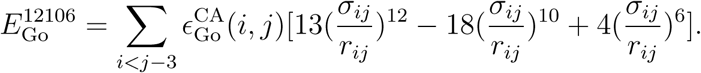

They represent the potential functions used in two popular versions of the coarse-grained Gō model: the Clementi-Onuchic model^29^ and the Karanicolas-Brooks model.^30^ The
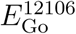
function is a modified LJ potential (Fig. S1) that incorporates a low energy barrier (a desolvation penalty) which has been shown to be able to increase the folding cooperativity of two-state folders,^88,89^ and improve model prediction.^90^ In both formulations,
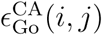
determines the depths of the potential wells and thus sets the strength of the native bias relative to the *E*_Profasi_ term. To keep the different models self-consistent, we built the coarse-grained native contact map used in the ProfasiGo model from the all-atom geometric occlusion contact map used in the pure Gō model described above. In particular, we considered two residues to be in contact in the ProfasiGo model if they share at least one atomic contact in the atom-based Gō model.

### MC simulations with Profasi and ProfasiGo model

The MC simulations with Profasi model and ProfasiGo model were performed by the modified versions of ProFASi^73^ or PHAISTOS software,^72^ both of which have implemented the Profasi force field.^75^ The patches for adding the Gō functions to the ProFASi and PHAISTOS software will be available at http://github.com/XXX.

We simulated *α*3W using parallel tempering/replica exchange (PT) with a set of eight temperatures ranging from 279K to 394K with the same interval. To get efficient sampling of the free energy landscape, we also used MUNINN, which employs the generalized multi-histogram equations^91,92^ and a nonuniform adaptive binning of the energy space, ensuring efficient scaling to large systems. We used a *β* (inverse temperature) ranging from 1.3 to 2.4, corresponding to a temperature range of 278K to 513K.

Three different elementary MC moves are used in the simulations: (a) biased Gaussian steps (BGS), (b) rotations of individual side-chain angles (Rot), (c) pivot-type rotations about individual backbone bonds (Pivot). The BGS move is semi-local and updates up to eight consecutive backbone degrees of freedom but keeps the ends of the segment approxi-mately fixed.

### MD simulations with pure Gō model

Simulations with the pure Gō model were performed with Gromacs 4.6.5.^93^ The dynamics of the systems was simulated using the Langevin thermostat with friction coefficient of *γ* = 1.0. Reduced units and a time step of 0.5 fs were used. Multiple trajectories were collected at a temperature range around the folding temperature (*T_f_*) for each protein system. The length of each simulation is 4×10^8^ simulation steps to include dozens of folding/unfolding transitions. We saved conformations every 2000 integration steps.

### Order parameters to characterise the folding mechanism

The fraction of native contacts, Q, has been shown to be good reaction coordinate in the study of protein folding.^32,94^ To describe the folding mechanism of the three-helical bundle proteins we also employed three local order parameters for helix formation, *Q*_H1_, *Q*_H2_, *Q*_H3_, and three order parameters that describe the pairwise assembly of helices, *Q*_H1-H2_, *Q*_H2-H3_, *Q*_H1-H3_. In all cases, *Q*_X_ is a measure of the progress of helix formation or assembly, by quantifying how far native contacting atoms are from their respective reference distances. More precisely, *Q*_X_ is a summation over the native contact pairs in the list denoted as X:

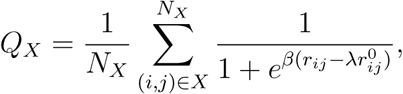

where X can be H1, H2, H3, H1-H2, H1-H3 and H2-H3, which are the lists of intra-segment contacts of helix1, helix2, helix3 and inter-segment contacts between them. Here, *r_ij_* is the distance between atom *i* and atom *j* in instantaneous structure (in units of nm), while
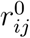
is the corresponding native distance. We set *β*=50 nm^−1^ and λ=1.5 in this work. Defined in this way, *Q_X_* values fluctuate between 0 (non-native) and 1 (native).

In addition, we employed four folding order parameters: *Q*_secondary_ to measure the fraction of native contacts within the three helices, *E*_HB_ (backbone hydrogen bond energy) to quantify the formation of secondary structure, *P*_helix_ to measure the proportion of *α* helical content, and *P*_beta_ to measure the proportion of *β* strand content, and two assembly order parameters: *Q*_tertiary_ to measure the fraction of native contacts between the three helices, and *E*_HP_ (hydrophobic energy), to quantify the formation of hydrophobic core.

### Selection of model proteins

Our goal was to develop the hybrid ProfasiGo model, and to test its range of applicability by benchmarking against other possible methods for studying protein folding. We thus focused our work on four three-helix bundle proteins whose folding mechanisms have previously been examined. The two proteins *α3*W and *α*3D are designed proteins that consist of three *α*-helices connected by two turns (Fig. 1A-B). According to the arrangement of the helices, the topology of *α*3W and *α*3D are considered to be left-handed and right-handed, respectively.^95,96^ In this sense they represent a pair of proteins with similar topologies but different conformational ‘chiralities’. We also chose the engrailed homeodomain (EnHD) and a designed thermostabilized homeodomain (UVF)^97^ (Fig. 1C-D) which are also three-helix bundle proteins. EnHD and UVF represent a pair of proteins with almost the same size and same handedness of the arrangement of the helices (Fig. 1 and Table 1). The similar topology of these four proteins allows us to use a consistent set of order parameters (as defined in the Method section) to characterise the folding mechanism. Because of the inclusion of a native-state bias, both the pure Gō model and the ProfasiGo model are expected to fold all four proteins to their native states. This is, however, not the case for the two physics-based force fields (Table 1). By looking across all four proteins, we can compare the folding mechanism observed in the ProfasiGo model with the results of three other models (a pure Gō model, the simple Profasi force field and an all-atom explicit solvent force field (Table 1).

**Figure 1:**
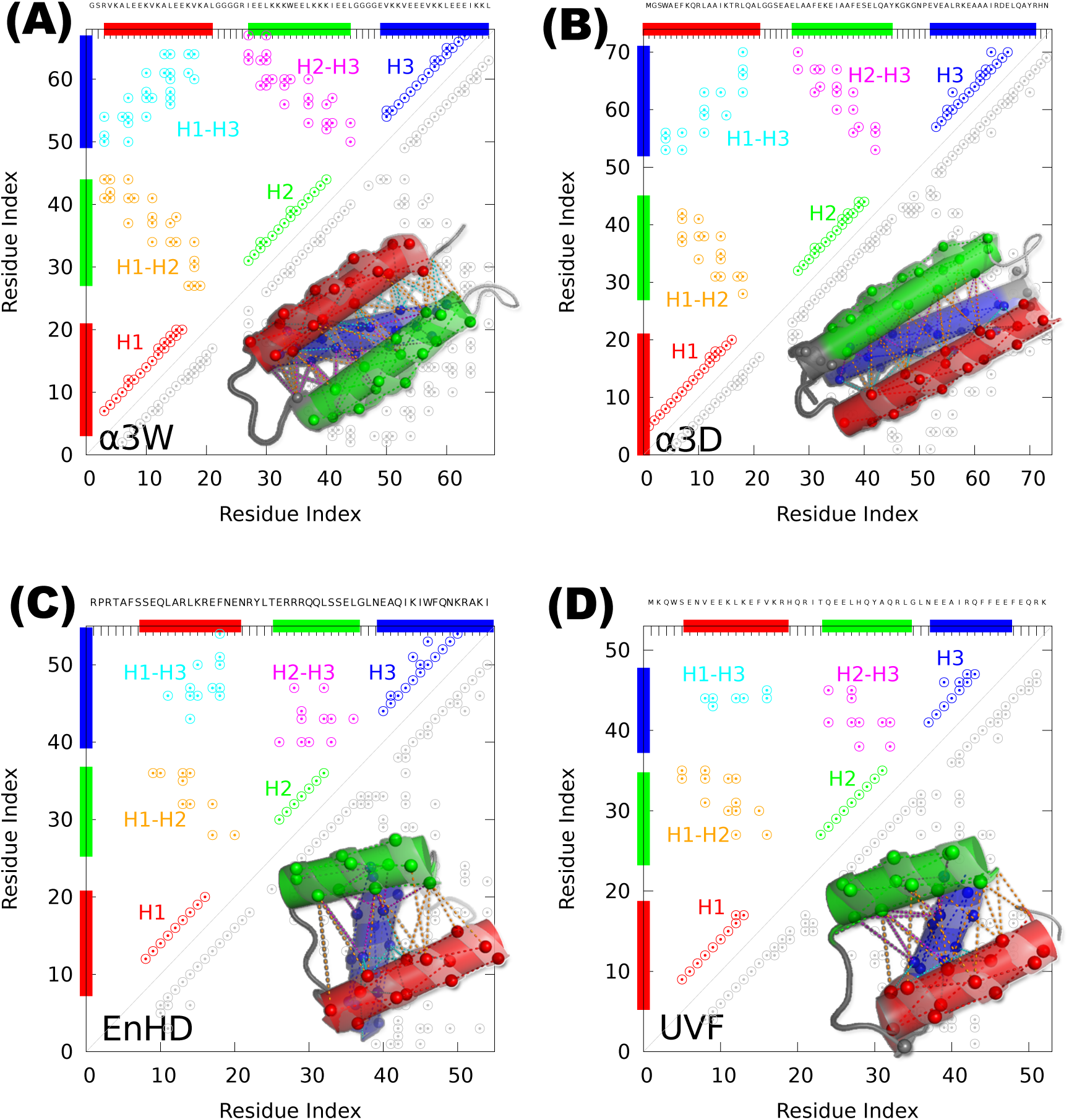
Contact maps and structure of the four three-helix bundle proteins studied here. Helices 1, 2 and 3 are coloured in red, green and blue, respectively. The inter-helix contacts between helix 1 and 2, between helix 1 and 3, and between helix 2 and 3 are coloured in orange, cyan and magenta, respectively. *α*3W and *α*3D share similar topology but different handedness of the orientation of the three helices; their sequence identity is 18%. EnHD and UVF have the same topology and handedness and a 23% sequence identity.

**Table 1:** Atomistic Models and their ability to fold *α*3W, *α*3D, EnHD and UVF

| System | Size (a.a.) | Topology | Profasi | SMOG | ProfasiGo | CHARMM22* |
| --- | --- | --- | --- | --- | --- | --- |
| $\alpha 3\text{W}$ | 67 | left-handed | Y | Y | Y | - |
| $\alpha 3\text{D}$ | 73 | right-handed | N | Y | Y | Y |
| EnHD | 54 | right-handed | - | Y | Y | N |
| UVF | 52 | right-handed | N | Y | Y | Y |

'Y', 'N' and '-' refer to the ability of the force field or model to fold the protein. 'Y': folded successfully, 'N': failed to fold, '-': unknown.

The remainder of the manuscript is thus constructed as follows. (i) We first compare the folding mechanism of a single protein (*α*3W) in the hybrid ProfasiGo model with that of the parent Profasi model. (ii) We then compare the folding mechanism of the four proteins under the ProfasiGo model to the results in a pure Gō model, and examine the dependency of the results on model parameters. (iii) Finally, we compare the folding mechanism of *α*3D and UVF in ProfasiGo and the all-atom, explicit solvent CHARMM22^∗^ force field simulations.

## Results and Discussion

### ProfasiGo and unbiased Profasi capture similar folding mechanisms of *α*3W

The designed *α*3W protein has previously successfully been folded by simulations with both coarse-grained models^98–100^ and all-atom force fields.^5,95,101–103^ In particular, its folding thermodynamics has been characterized by Irbäck et al. using the pure Profasi force field,^75^ making it particularly suitable to be used for testing and calibrating our ProfasiGo model.

We sampled the free energy landscape of *α*3W with the ProfasiGo model in the multicanonical (’flat-histogram’) ensemble using the MUNINN software.^91,92^ Such a generalized ensemble method can not only improve sampling efficiency, but also directly helps determine the melting temperature where folding and unfolding transitions typically occur most frequently, a time-consuming but often necessary process in MD or MC simulations.^37,64^ Subsequently, the thermodynamics properties at any temperature of interest can be obtained by reweighting techniques.^104^

Throughout this manuscript we explore protein folding through such enhanced sampling simulations, examining folding mechanisms by analysing and comparing free energy profiles. As an example, we calculated three order parameters for folding, *Q*_H1_, *Q*_H2_ and *Q*_H3_ (see definitions in Models and Methods), that describe the formation of each of the three helices, and project the conformational space onto the two-dimensional free energy surfaces spanned by combinations of these coordinates (Fig. 2A). Free energy surfaces as a function of such order parameters have been widely used to elucidate protein folding and assembly mechanisms through minimum free energy pathways.^62,64^ For *α*3W, we observe that there are no low energy pathways along the diagonal lines in these free energy surfaces (F(*Q*_H1_,*Q*_H2_), F(*Q*_H1_, *Q*_H3_) and F(*Q*_H2_,*Q*_H3_)), indicating that the folding of the three helices is independent of one another, without strong coupling. Furthermore, there are multiple possible folding routes, suggesting heterogeneity in the order of formation of the three helices.

**Figure 2:**
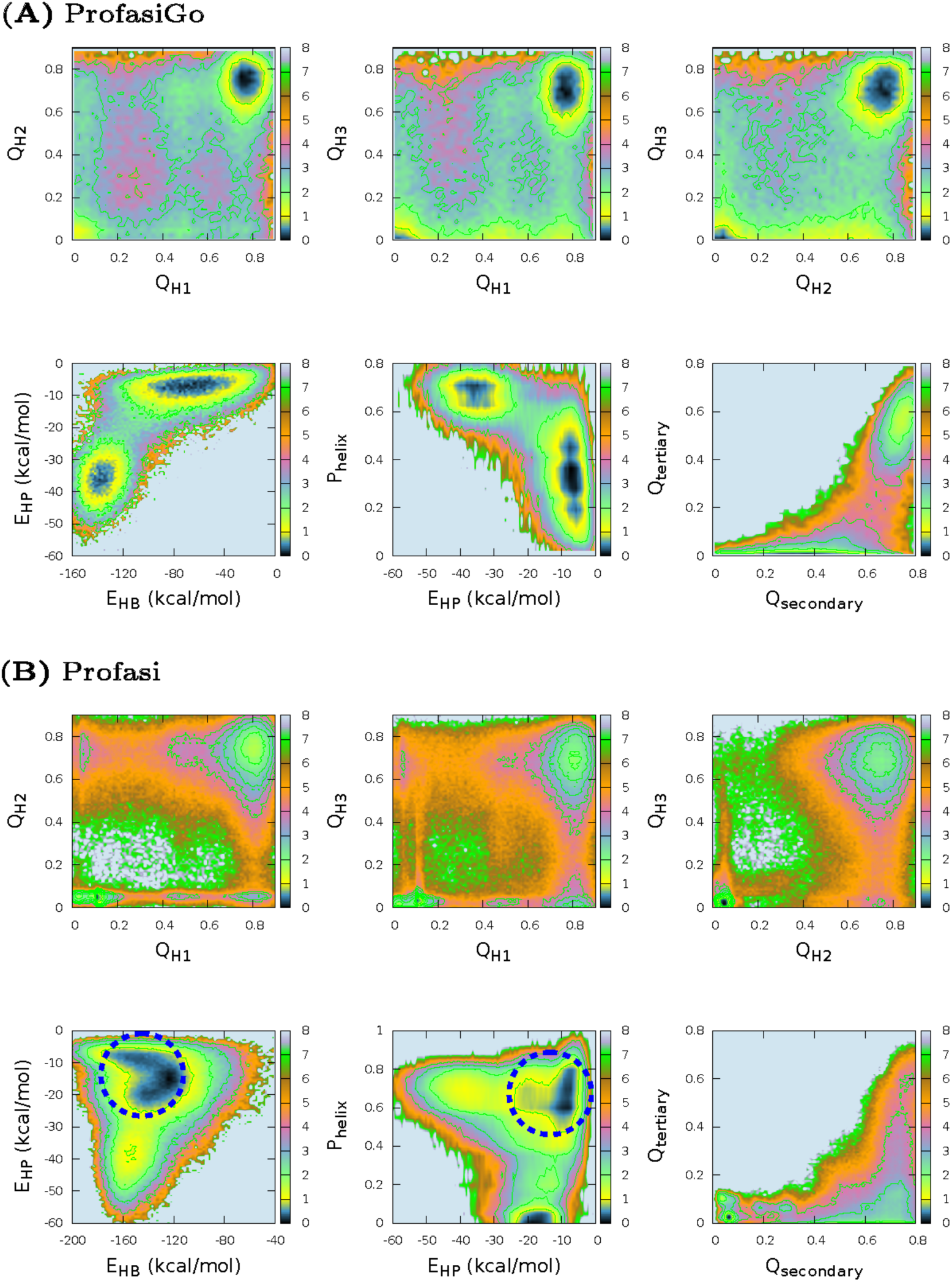
Similar folding mechanisms for *α*3W in the Profasi and ProfasiGo models. (A) Free energy surfaces from MC simulations with the ProfasiGo model with *ϵ*_Go_=0.2 and reweighted to 346K. The top three panels show F(*Q*_H1_,*Q*_H2_), F(*Q*_H1_,*Q*_H3_), F(*Q*_H2_,*Q*_H3_), where *Q*_H1_, *Q*_H2_ and *Q*_H3_ are the fraction of native intra-helical contacts. The bottom three panels show F(*E*_HB_,*E*_HP_), F(*E*_HP_,*P*_heiix_) and F(*Q*_secondary_,*Q*_tertiary_). Here *E*_HB_ and *E*_HP_ are the backbone hydrogen bond energy and hydrophobic energy, respectively, while *P*_helix_ is fraction of helix formed. Low free-energy pathways are labeled by white arrows. (B) Same plots as in A, but from MC simulations by the Profasi model and analyzed at T=303K. Possible misfolded states are highlighted by blue dashed circles. All free energies are in units of kcal/mol.

In addition to examining the order of formation of the different secondary structure elements, we also analysed the relationship between formation of secondary and tertiary structure; such an analysis would be useful to distinguish between a nucleation condensation model, diffusion collision or framework model, or hydrophobic collapse model for folding. We thus calculated two additional local order parameters: *Q*_secondary_ to measure the fraction of native contacts within the three helices and *P*_helix_ to measure the fraction of helical content, and two order parameters aimed to capture tertiary interactions: *Q*_tertiary_ to measure the fraction of native contacts between the three helices and the hydrophobic energy, *E*_HP_, to quantify the energy of forming the hydrophobic core. The two-dimensional free energy surfaces as a function of *E*_HP_ and *P*_helix_, and *Q*_secondary_ and *Q*_tertiary_ illustrate a clear pathway that helix folding occurs before the formation of the hydrophobic core (Fig. 2A). Therefore, the thermodynamic free energy analysis suggests the folding of *α*3W in this model can be well described by the diffusion collision model^105^ by which the native secondary structures are formed before the tertiary structures. This conclusion is consistent with previous studies.^66,101^

Having analysed the folding free energy surfaces in our new hybrid model we proceed to compare with the surfaces obtained in the same model, but without the native bias. The basic hypothesis is that, for the proteins that can be folded by the physics-based (non-Gō) Profasi model, the hybrid model would retain most of the features observed in the less biased model. By taking the same protocol as previously applied in the work of Irbäck et al,^75^ we performed replica-exchange MC simulations with the pure Profasi model, and determined the folding free energy surfaces for *α*3W (Fig. 2B). By comparing with the free energy surfaces obtained from the hybrid ProfasiGo model (Fig. 2A), we find overall similar shapes of the free energy landcapes, suggesting a similar mechanism for folding of *α*3W. In addition to additional ‘roughness’ of the landscape in the pure Profasi model, the major difference are two intermediate states present on the free energy surface F(*E*_HP_,*P*_helix_) sampled by the Profasi model. Inspection of the structures of these intermediate states revealed that the consist of very long helices or a high proportion of beta strands, but without any substantial hydrophobic packing, which we consider to be artefacts of the Profasi model.

In summary, the results suggest that *α*3W folds by a diffusion collision mechanism in both the pure Profasi model and in the hybrid ProfasiGo model. This observation supports the idea that the introduction of the native-biased Gō potential mostly acts to smoothen the energy landscape of protein folding, but does not substantially change the folding mechanism. This in turn suggests that simulations of protein folding with the hybrid model would yield realistic folding mechanisms even in cases where folding simulations are not possible with the pure Profasi model.

### Comparing the hybrid model with a pure Gō model

While the ProfasiGo model shows the ability to reproduce the folding mechanism as revealed by the unbiased Profasi model, we also examined whether the pure Gō model would suggest a similar mechanism. We performed constant temperature MD simulations using a pure all-atom Gō model generated by SMOG server with default parameters. By performing MD simulations on different temperatures ranging from 100 to 130K, which cover the expected folding temperature for normal proteins,^35^ we found the folding temperature for *α*3W in SMOG Gō model to be around 109K. We then projected the MD trajectories onto the same folding order parameters as we used in the ProfasiGo model (Fig. S2). Unexpectedly, the results are rather different, and indicate a more strongly coupled folding process for all three helices. Thus the results from the pure Gō model suggest a nucleation condensation mechanism, in contrast to the diffusion collision mechanism revealed by both the Profasi and ProfasiGo models. Without more detailed experimental data available for the folding of *α*3W it is difficult to know which model is more realistic, but our hypothesis is that the combination of the physical and Gō model in principle provides access to more complex and varied mechanisms.

### Distinguishing between folding mechanisms of *α*3W and *α*3D

Next, we studied the folding mechanism of *α*3D, which has similar topology but different handedness of the packing the three helices. We carried out MC simulations of *α*3D with the same strategy as for *α*3W, and compared the resulting free energy surfaces of the two models. The results suggest significant difference in the free energy surfaces between *α*3W and *α*3D (Fig. 3). Not only is the folding pathway of the three helices in *α*3D distinct from *α*3W, but the order of the formation of secondary and tertiary structure is also remarkably different. For *α*3W, the results suggest that the secondary structure forms before the hydrophobic core, while for *α*3D, the formation of secondary structure and the hydrophobic core are strongly coupled. Therefore, the folding mechanisms of *α*3W and *α*3D represent two different models: diffusion collision and nucleation condensation, respectively. This conclusion is consistent with recent work carried out by Shao with integrated-tempering-sampling MD simulations.^101^

**Figure 3:**
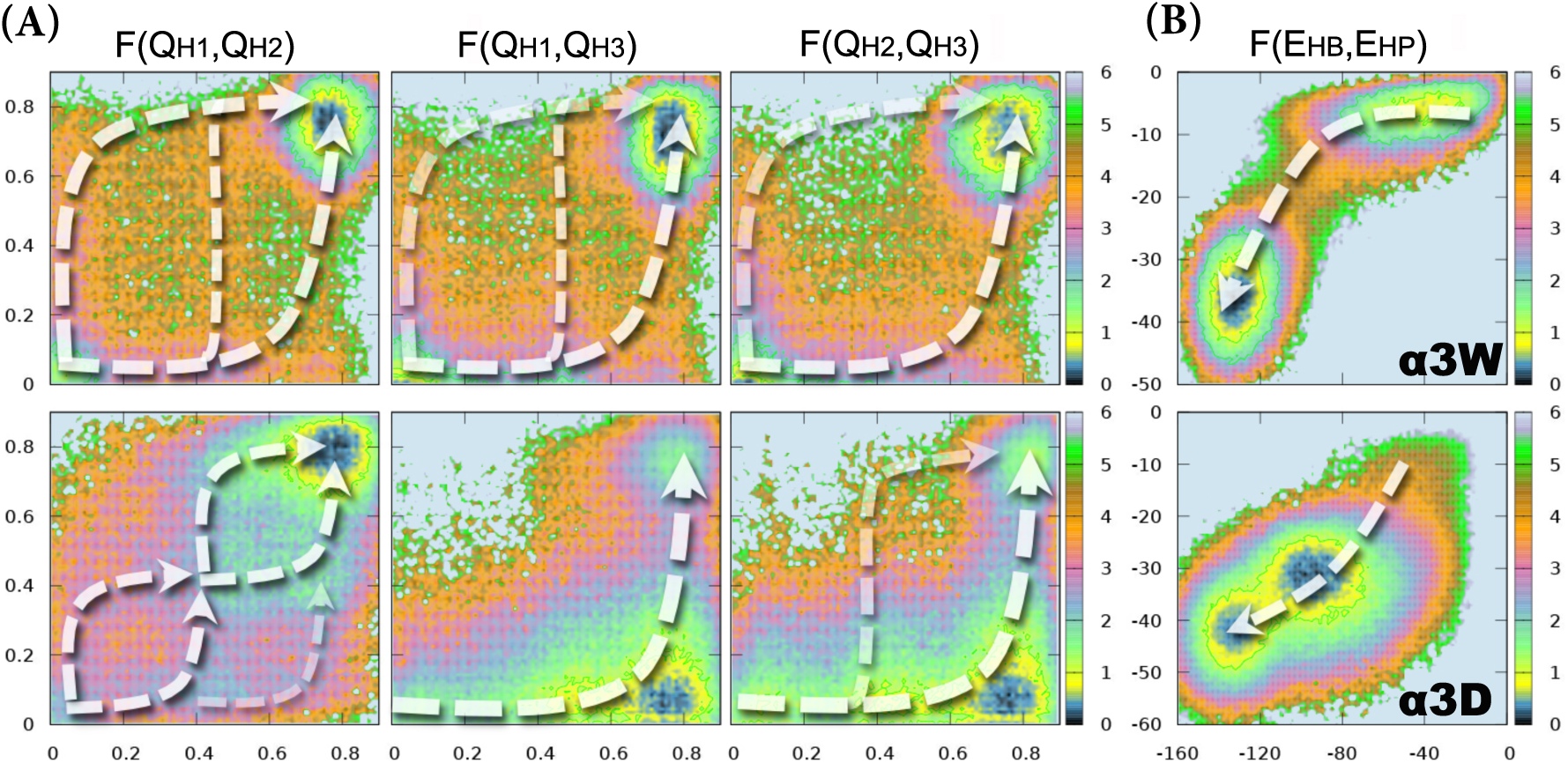
Distinct folding mechanism of *α*3W and *α*3D suggested by the ProfasiGo model. A) F(*Q*_H1_, *Q*_H2_), F(*Q*_H1_, *Q*_H3_) and F(*Q*_H1_, *Q*_H2_). (B) F(*E*_HB_, *E*_HP_). The results for *α*3W and *α*3D are shown in the first and second row, respectively. Their free energy landscapes were sampled by MUNINN with *ϵ*_GO_ = 0.3, and reweighted at *T_f_*=373K for *α*3W and *T_f_*=339K for *α*3D, respectively. Low free-energy pathways are labeled by white arrows. All free energies are in units of kcal/mol.

### Distinguishing between folding mechanisms of EnHD and UVF

After demonstrating that the ProfasiGo model can distinguish the folding mechanism of a pair of proteins with similar topology but different handedness of the packing, we proceeded to a more challenging case: to capture the differences in the folding mechanism of a pair of proteins with the same topology; such differences are generally difficult to capture within a pure Gō model.^30^

We chose the engrailed homeodomain (EnHD) and its thermostabilized variant (UVF)^97^ as our target systems (Fig. 1C-D and Table 1), and compared the global free energy landscape by projecting the conformational space onto a few global order parameters, including *E*_HB_ (backbone hydrogen bond energy) and *E*_HP_ (hydrophobic energy). The free energy surfaces are quite different between EnHD and UVF (Fig. 4), suggesting the presence of folding intermediate states, which previously have been proposed by both simulation and experimental studies.^106,107^ The two-dimensional free energy surfaces of F(*E*_HB_,*E*_HP_) suggest that EnHD has a tendency to form secondary structure before the formation of hydrophobic core, while UVF tends to form the hydrophobic core coupled with the formation of secondary structures. In addition, the conformational distribution in the free energy surfaces suggests that UVF can sample conformational regions with lower hydrophobic energy. This may be explained by the fact that UVF has higher percentage of hydrophobic residues.^101^ In any case, our results suggest that the global folding mechanism of the two proteins can be distinguished by the ProfasiGo model despite the fact that they share almost exactly the same topology.

**Figure 4:**
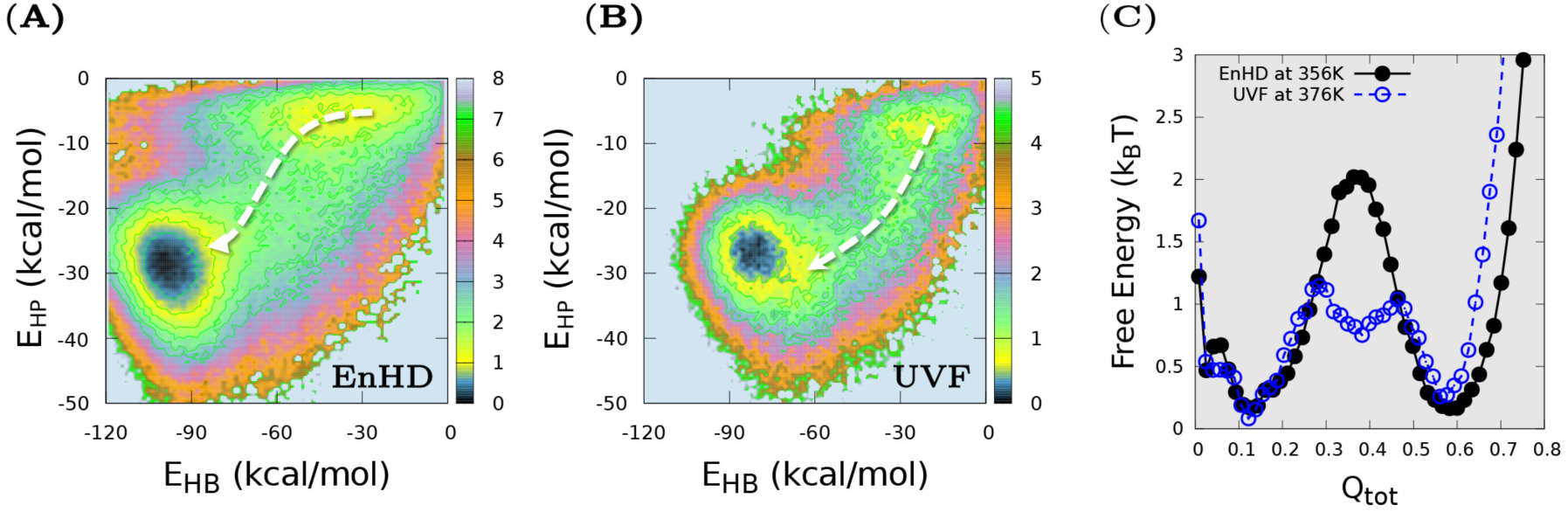
The ProfasiGo model suggests distinct free energy profiles for two proteins with the same topology (EnHD and UVF). (A-B) F(*E*_HB_,*E*_HP_) for EnHD and UVF, respectively. (C) F(*Q*_tot_) for EnHD and UVF. The results are from multicanonical MC simulations with the ProfasiGo model and reweighted to their corresponding folding temperatures (356 K and 376 K for EnHD and UVF, respectively). Low free-energy pathways are labeled by white arrows.

### Effects of the strength and shape of the Gō potential

The strength of the Gō potential relative to the physical potential, determined by *ϵ*_Go_, is a free parameter in the ProfasiGo model. To assess how sensitive the results are to the choice of this value, we performed MC simulations of *α*3W with the same generalized ensemble method but different values for *ϵ*_Go_. The resulting free energy landscapes at their corresponding *T_f_* show very similar free energy surfaces for different Gō strengths, indicating the same folding mechanism. Thus, our results suggest the folding mechanism predicted by the ProfasiGo model is quite robust to the variety of Gō strength with this range. This conclusion is also supported by comparison of the free energy surfaces projected onto other order parameters, and by the corresponding simulations on UVF (Fig. S3). Unsurprisingly, we find that the free energy surfaces with different *ϵ*_Go_, e.g. as a function of total potential energy (*E*_tot_) and RMSD, show that the energy landscapes become more ‘funnelled’ as the strength of Gō potential increases (Fig. 5 and Fig. S3).

**Figure 5:**
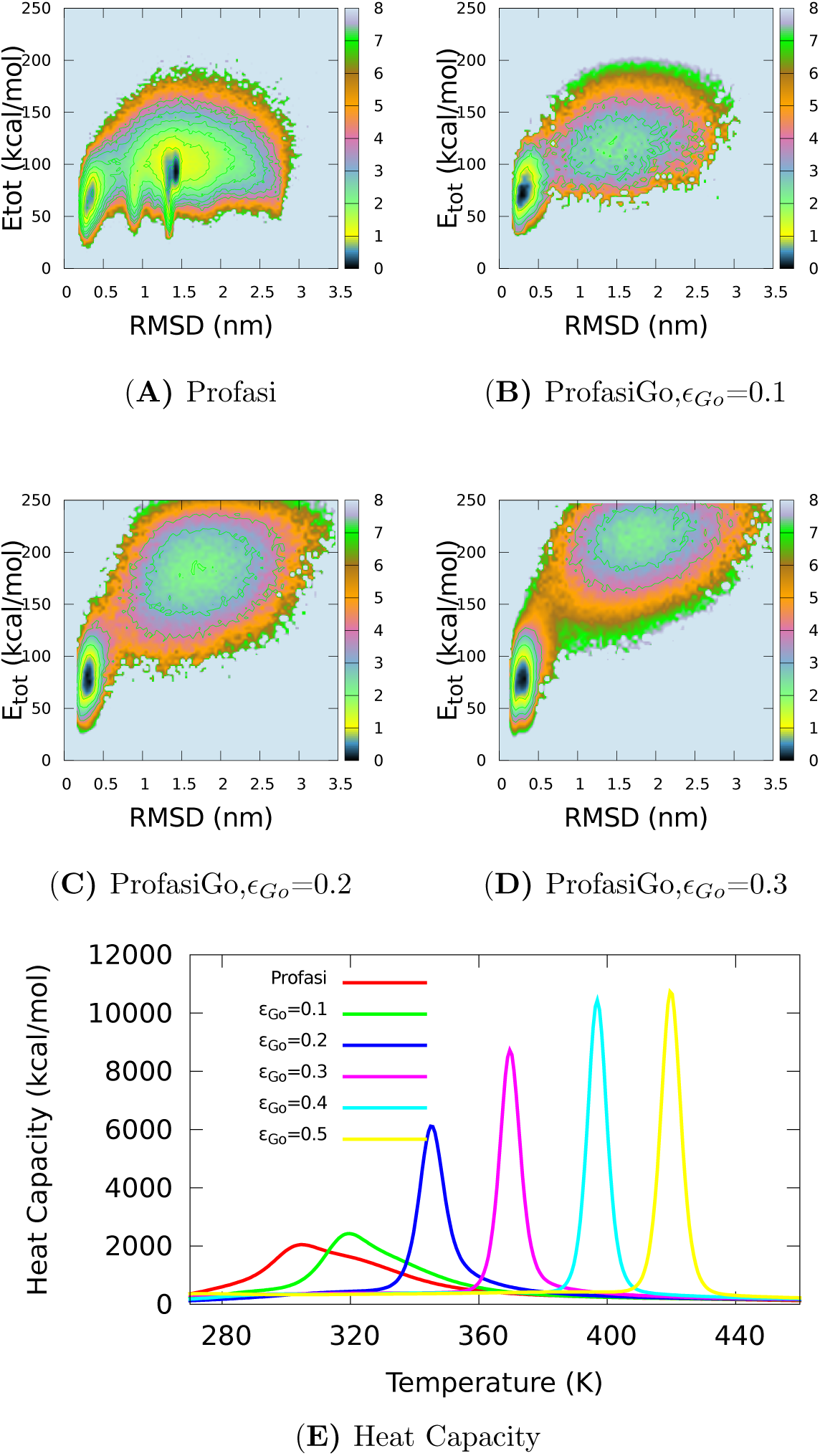
The Gō potential makes the energy landscape more funnelled. (A) Profasi model of o3W (equal to a ProfasiGo model with *ϵ*_Go_ = 0.0) at *T_f_* = 303*K*; (B) ProfasiGo model with *ϵ*_Go_ = 0.1 at *T_f_* = 315*K*; (C) ProfasiGo model with *ϵ*_Go_ = 0.2 at *T_f_* = 346*K*; (D) ProfasiGo model with *ϵ*_Go_ = 0.3 at *T_f_* = 370*K*. (E) The curves of heat capacity in ProfasiGo models with variant *ϵ*_Go_ ranging from 0.0 to 0.5.

We also tested two types of Gō potential (12-10 and 12-10-6 forms as described in the Methods section), again using *α*3W as the test case, and found that the folding mechanism predicted by the ProfasiGo model is not sensitive to the shape of Gō potential, though we did find a small increase of the free energy barrier (Fig. S4) when using the 12-106 form. Previous simulations based on the coarse-grained Gō models have shown that the introduction of the desolvation barrier can help rationalise the diversity in the protein folding rates as well as the experimentally observed folding cooperativity.^88,89,108^ Our observation that the mechanism is less sensitive to the choice of the functional form in the context of the ProfasiGo model implies that the non-native interactions are fairly well captured by the physics-based term, thus alleviating this responsibility from the structure-based term.

### Comparison of ProfasiGo, Gō and all-atom MD simulations

The availability of long time-scale, unbiased MD simulations of both *α*3D and UVF performed with ANTON^5^ allowed us to conduct a final experiment, comparing the free energy landscapes and protein folding mechanisms obtained by different models, spanning from a pure Gō model (SMOG), the hybrid ProfasiGo model, to an explicit solvent, all-atom force field (CHARMM22^∗109^ with TIP3P water^110^).

To examine the folding mechanisms obtained from different models, we projected the folding trajectories of *α*3D and UVF onto the two-dimensional free energy surfaces arising as combinations of *Q*_H1_, *Q*_H2_ and *Q*_H3_ (Fig. S5 and Fig. S6). The results suggest that the three helices fold independently in the ProfasiGo model, while they are strongly coupled in the pure Gō model. The folding of the three helices is more complex in CHARMM22^∗^, but is consistent with ProfasiGo in the sense that it also finds the helices to form independently.

To examine the global free energy landscape, we further projected the folding trajectories onto two-dimensional free energy surfaces of F(*Q*_secondary_, *Q*_tertiary_) for *α*3D (Fig. 6) and UVF (Fig. S7). We find that the free energy landscape from the ProfasiGo model is more similar to those of CHARMM22^∗^ than those from the pure Gō model.

**Figure 6:**
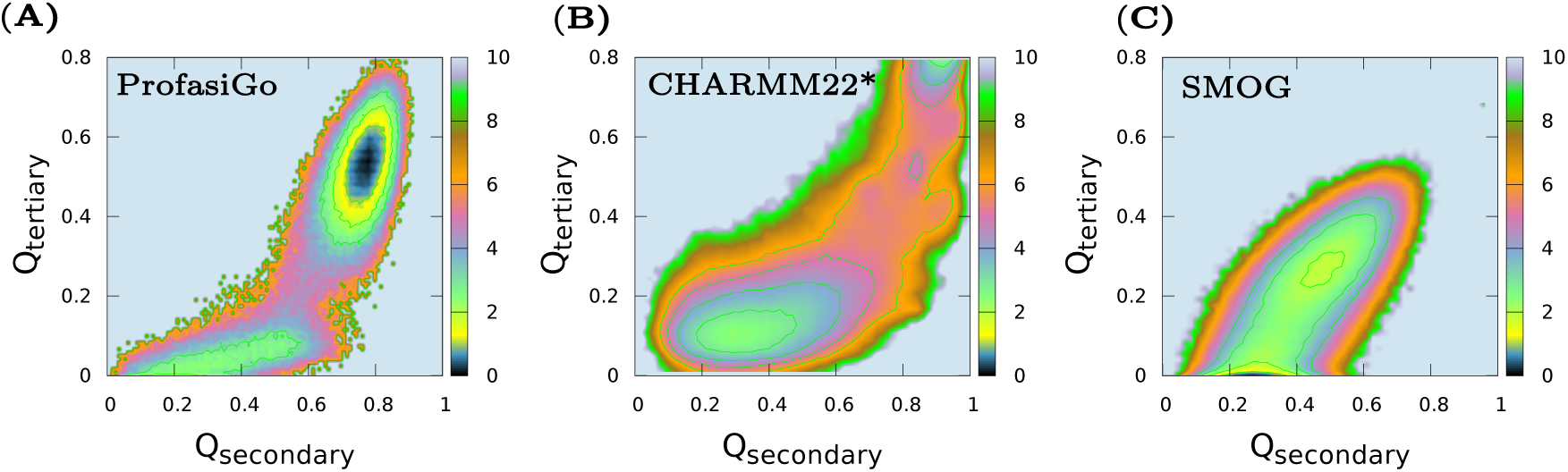
Comparison of the global free energy landscapes of *α*3D obtained from a pure Gō model, the hybrid ProfasiGo model and an explicit solvent force field. The results show the free energy surfaces of *α*3D as a function of *Q*_Secondary_ (the fraction of native contacts within the three helices) and *Q*_tertiary_ (the fraction of native contacts between the three helices) obtained from the ProfasiGo model with *ϵ*_Go_ = 0.5 (A), SMOG (B) and CHARMM22^∗^ (C). All free energies are in units of kcal/mol.

## Conclusions

Here we introduce a model that serves as a hybrid between an atomistic physics-based potential and a residue-level structure-based (Gō) model, in the context of a Monte Carlo simulation framework. We demonstrate that our model has the ability to successfully capture the protein folding mechanism at a level similar to a pure physics-based model. The model provides features not available with traditional structure-based approaches; for example, it is capable of distinguishing between different folding pathways for topologically similar proteins. Finally, the procedure is complementary to physics based models in cases where these fail to fold to the native state (or do so excessively slowly).

Our procedure has a some limitations. Like for most force fields, the experimental folding temperature cannot be perfectly reproduced, and the folding temperature must therefore be located by scanning a range of temperatures. Secondly, the fact that we have chosen Monte Carlo as the basis for our approach makes it difficult to obtain realistic kinetic information. This could potentially be be mitigated by careful selection of moves, or restricting the analysis to longer time-scales.^111,112^

The Monte Carlo approach, however, provides substantial benefits in terms of computational efficiency. Our procedure requires only a few weeks of computation on a single CPU to obtain converged simulations on modest size proteins (with 40-80 residues), a dramatic improvement over comparable explicit solvent force field simulations. This makes it attractive for rapidly probing structure-mechanism relationships, taking input either directly from native structures or from indirect structural information derived from e.g. NMR spectroscopy or co-evolutionary analysis.^77,113^ The ProfasiGo model may also serve as an efficient atomistic model for sampling conformational space of large proteins, which can be refined a posteriori by reweighting with available experimental data.^114,115^ Indeed, while we have here used the ProfasiGo model in the context of protein folding, we also expect it to be useful in providing access to conformational dynamics within folded states.

Overall, our results suggest that the ProfasiGo model can serve as a useful middle ground that combines the simplicity and efficiency of the Go-type models and the accuracy and high computational cost of explicit solvent MD simulations.

## Acknowledgement

K.L.-L. acknowledges funding by a Hallas-Møller Stipend from the Novo Nordisk Foundation and the BRAINSTRUC initiative from the Lundbeck Foundation. W.B. acknowledges funding from VILLUMFONDEN.

## Supporting Information Available

Figure S1, Two popular functional forms of Gō potentials. Figure S2, A standard Gō model suggests a different folding mechanism of *α*3W compared to the Profasi and ProfasiGo models. Figure S3, Evidence that the Gō potential can funnel the free energy landscape by reducing the population of misfolded states of UVF. Figure S4, Predicted folding mechanism by the ProfasiGo model is not sensitive to the mathematical form of the Go potential in the case of α3W. Figure S5, Comparison of the folding mechanisms of *α*3D obtained from a pure Gō model, a hybrid ProfasiGo model and an explicit solvent force field. Figure S6, Comparison of the folding mechanism of UVF obtained from a pure Gō model, a hybrid ProfasiGo model and an explicit solvent force field. Figure S7, Comparison of the global free energy landscapes of UVF obtained from a pure Gō model, a hybrid ProfasiGo model and an explicit solvent force field. This material is available free of charge via the Internet at http://pubs.acs.org.

This material is available free of charge via the Internet at http://pubs.acs.org/.

## Graphical TOC Entry

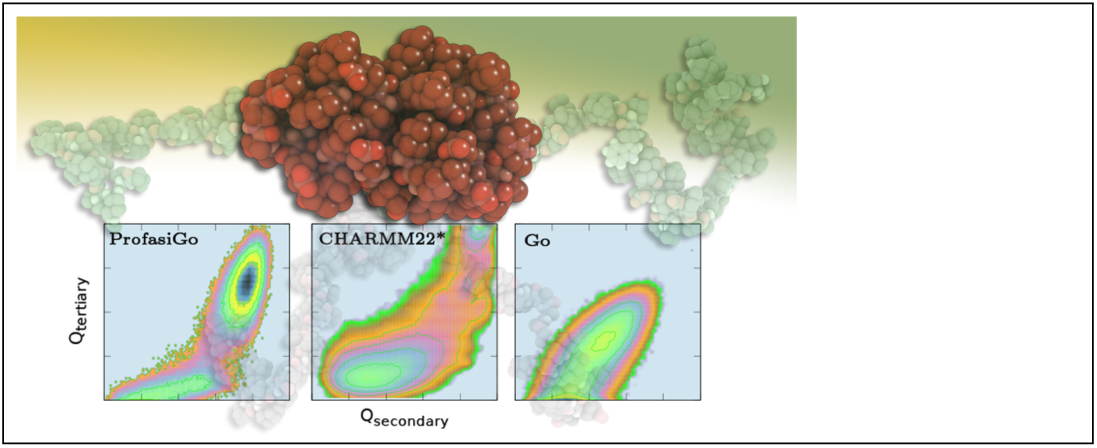

**Figure S1:**
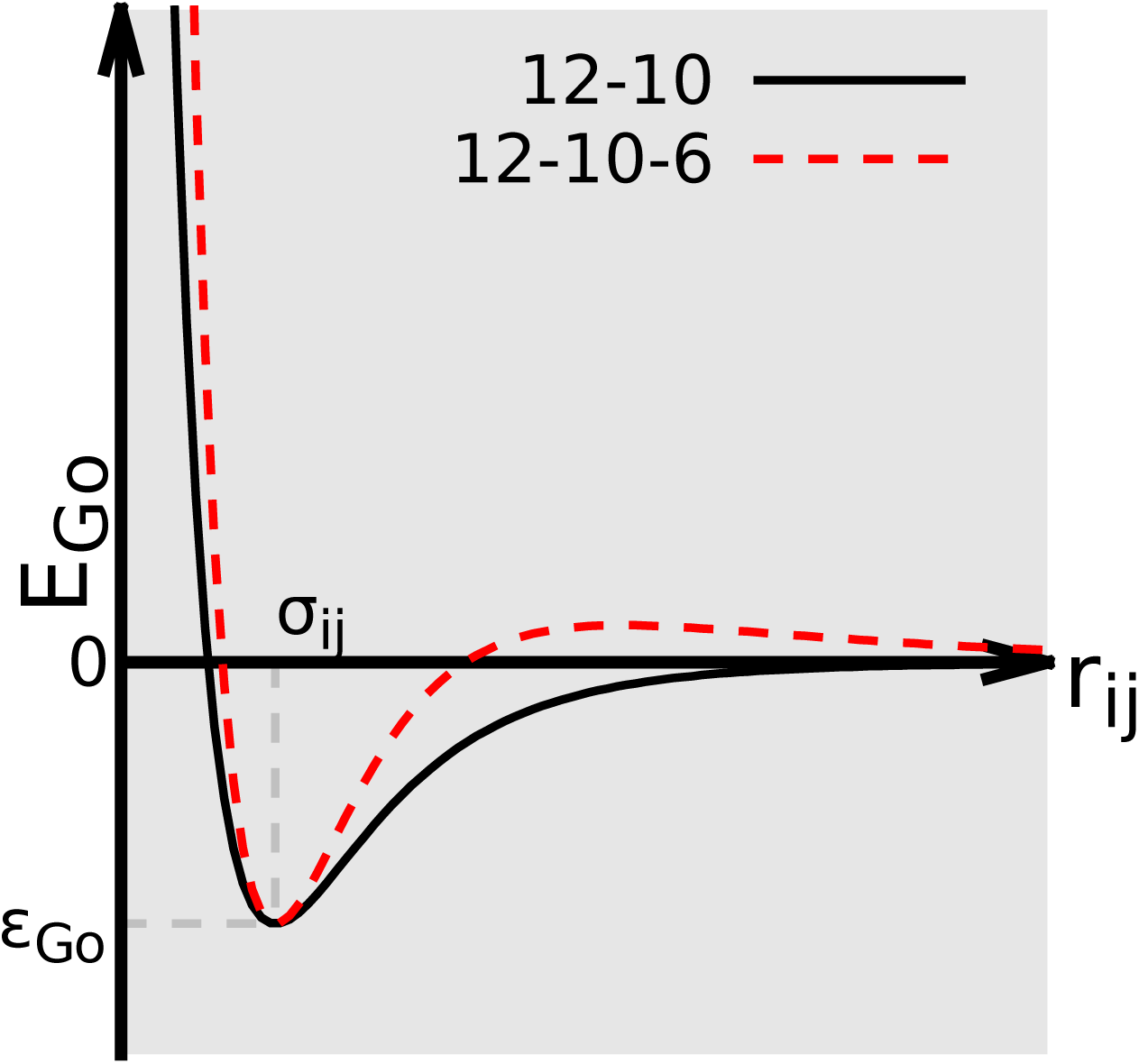
Two popular functional forms of the Gō potential. The two curves show the 12-10 Lennard-Jones-like potential, and the modified 12-10-6 potential, as black solid and red dashed lines, respectively. The 12-10-6 potential has a low energy barrier designed to mimic the desolvation effect.

**Figure S2:**
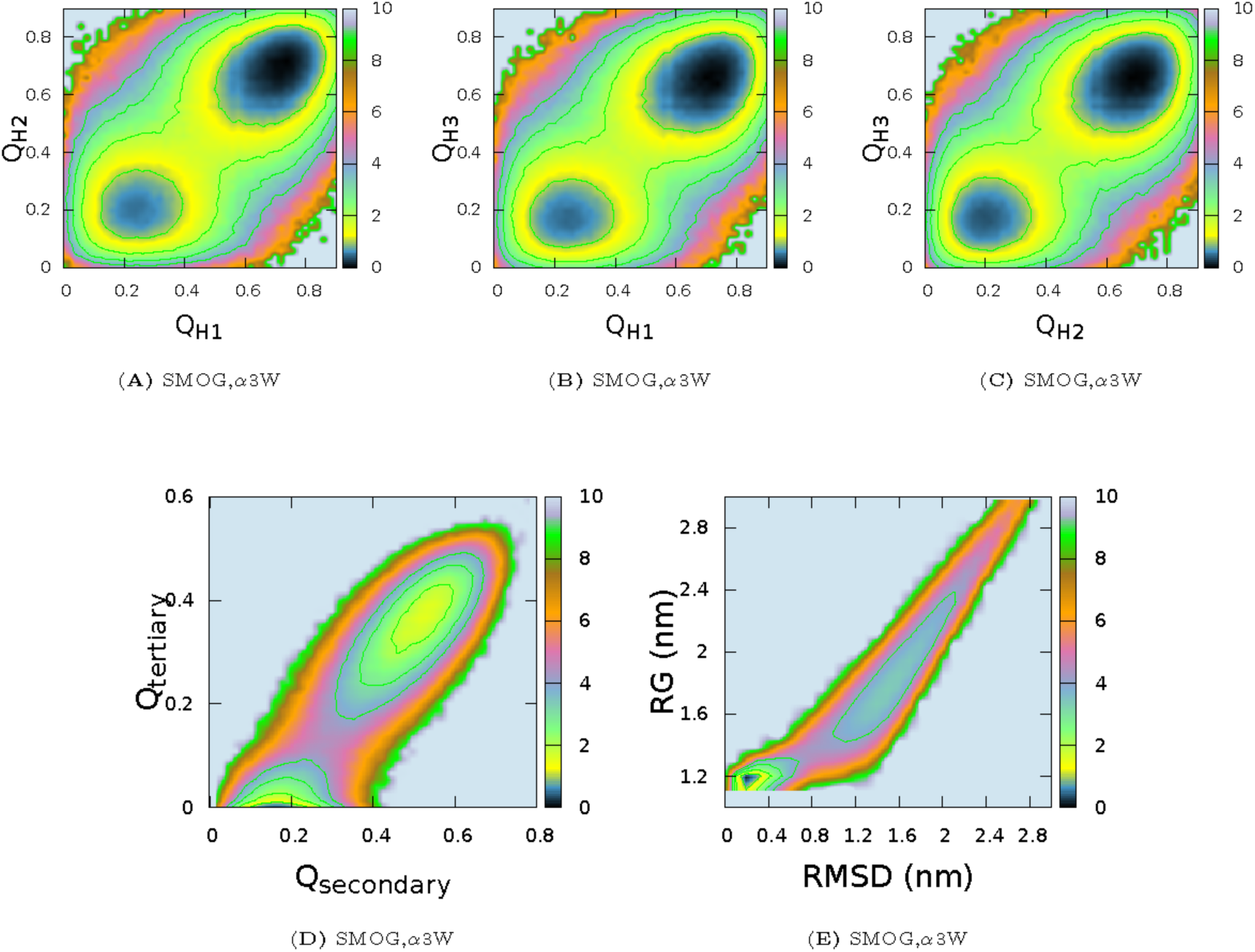
Free energy landscape of *α*3W in the pure Gō model. (A) F(*Q*_H1_,*Q*_H2_). (B) F(*Q*_H1_,*Q*_H3_). (C) F(*Q*_H2_,*Q*_H3_). (D) F(*Q*_secondary_,*Q*_tertiary_). (E) F(RMSD,RG). Comparison with the Profasi and ProfasiGo models (Fig. 2 in the main text) reveals a substantially different mechanism.

### Increasing the ‘foldability’ of a physics-based force field by adding a native bias

The free energy landscape, F(RMSD,*E*_tot_), in simulations of UVF with *ϵ*_GO_=0.3 reveals a non-native state (RMSD=10.0) with a comparable free energy and internal energy as the native state (Fig. S3A). Note that these results were based on *ϵ*_GO_=0.3 (Fig. S3A). In this case, increasing the native bias (*ϵ*_GO_=0.4) substantially decreases the stability of this state (Fig. S3B). Similar results were obtained for *α*3W, when comparing the free energy landscape in the absence of the native bias with those of the ProfasiGo model (Fig. 5 in the main text).

There are two key ingredients that help determine the whether an energy landscape is ‘well funnelled’: the energy gap between the native and nonnative state and the energetic fluctuations in the non-native states.^1^ The former determines the steepness of the energy funnel, while the later controls the roughness of the energy funnel. Maximization of their ratio has been used to guide the optimization of the force field parameters by following the minimal frustration principle and maximize the ‘funnelledness’ of the protein energy landscape.^2^ Here, we provided evidence that the introduction of native structure-based information into a physics-based force field can funnel the energy landscape not only by increasing the steepness (native state becomes more energetically favourable), but also by decreasing the roughness (the energy fluctuation or the width of the energy distribution of the non-native states becomes more narrow).

The results in Fig. S3 also suggests that the pure Profasi model (equivalent to ProfasiGo with *ϵ*_GO_=0) will not have the correct native state of UVF as its free energy minimum, since the non-native state becomes progressively more populated as the native bias is decreased. We thus suggest that analyses such as these could be useful to identify force field problems in cases where sampling the folding landscape is prohibitively difficult.

**Figure S3:**
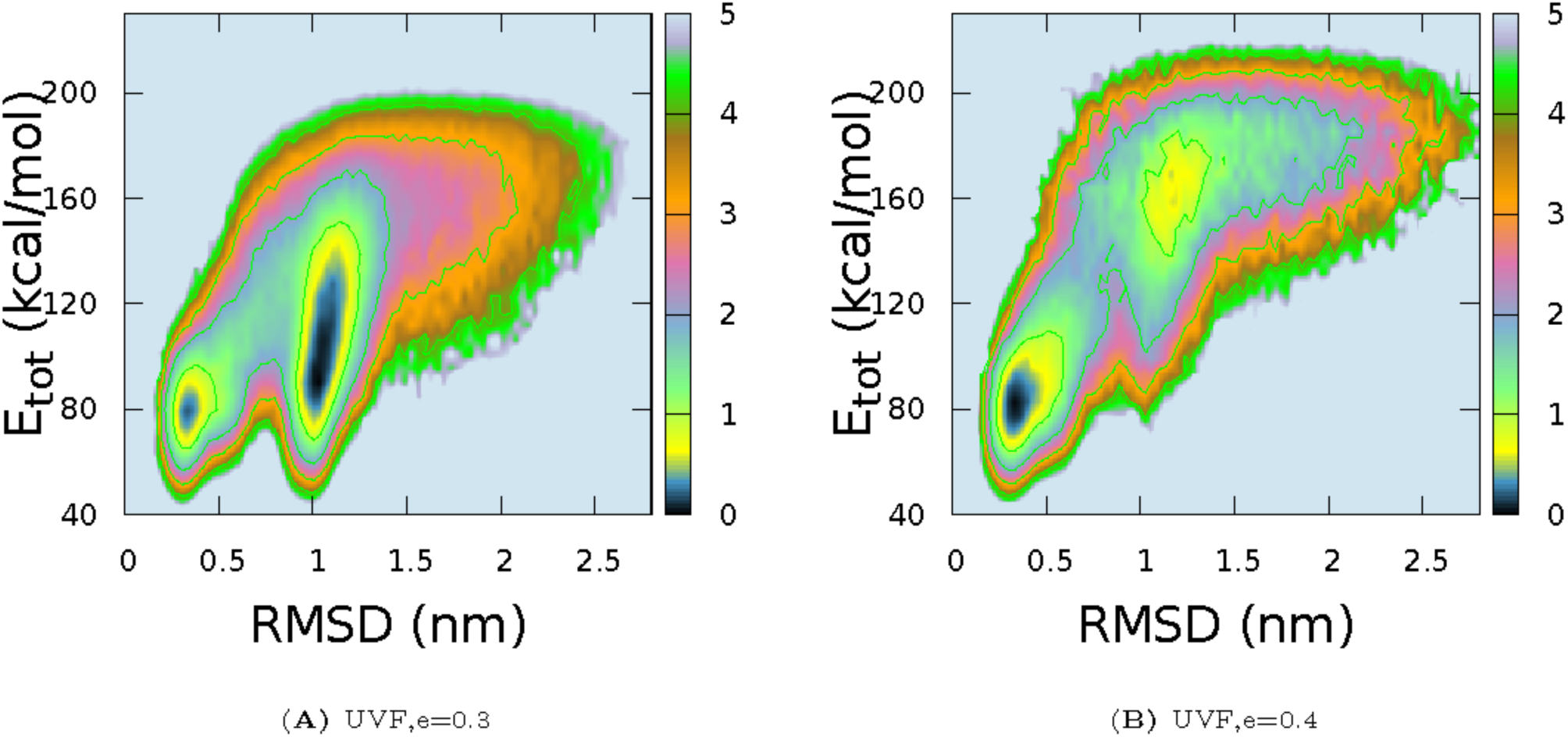
Evidence that Go potential can funnel the free energy landscape by reducing the population of misfolded states in the case of UVF. (A) Two-dimensional free energy surfaces as a function of RMSD and *E*_tot_ at *ϵ*_Go_=0.3. (B) Two-dimensional free energy surfaces as a function of RMSD and *E*_tot_ at *ϵ*_Go_=0.4. Note that the results were from multicanonical MC simulations by ProfasiGo model of UVF and reweighted to corresponding folding temperature.

**Figure S4:**
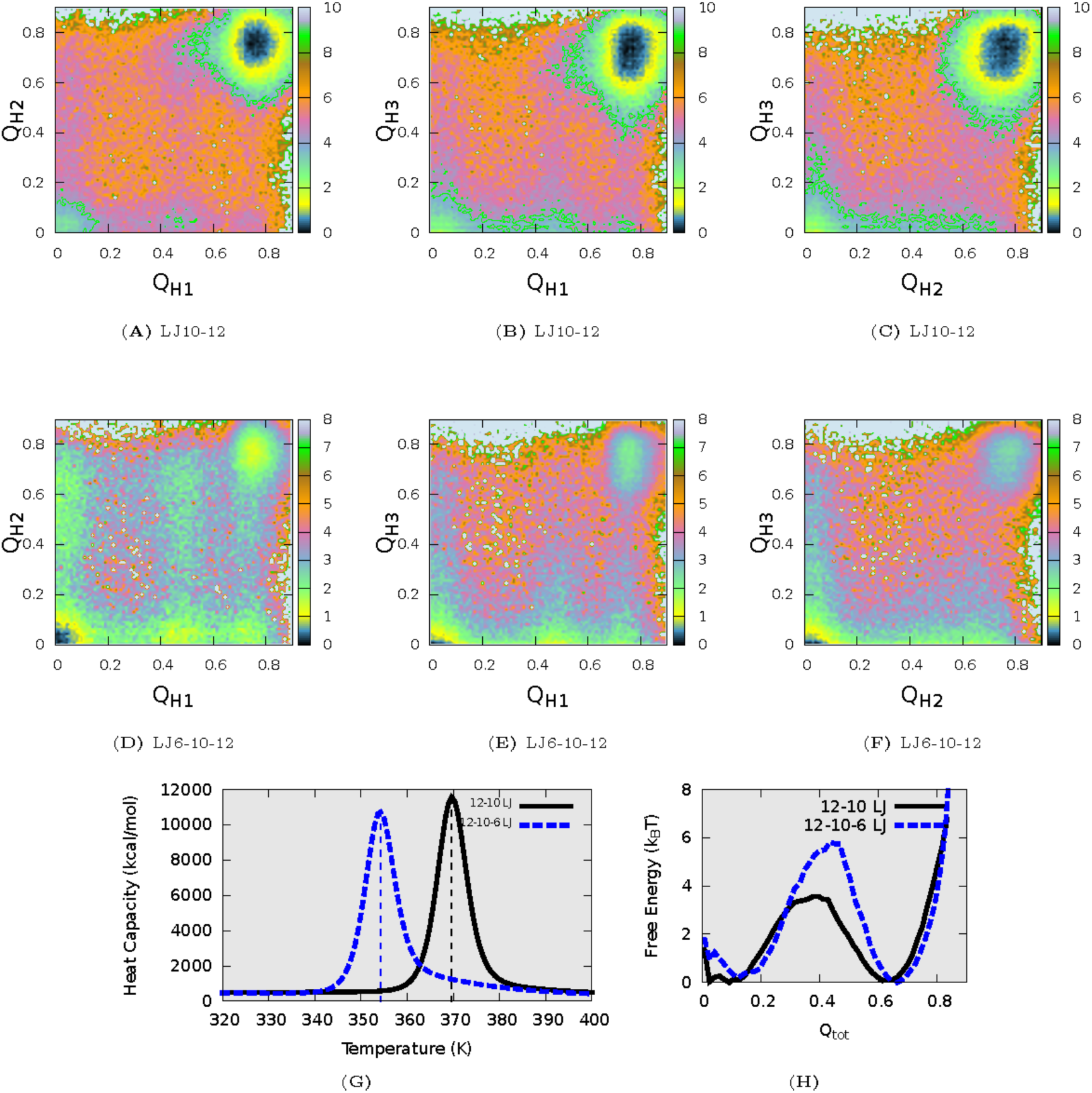
The free energy landscape in the hybrid ProfasiGo is not sensitive to the mathematical form of the Gō potential. (A–F) Free energy landscape of *α*3W using the 12-10 potential (A–D) or 12-10-6 potential. (A–C) F(*Q*_H1_, *Q*_H2_), F(*Q*_H1_, *Q*_H3_), F(*Q*_H1_, *Q*_H2_) and F(*Q*_all_) with the 12-10 potential. (D–F) F(*Q*_H1_, *Q*H_2_), F(*Q*_H1_, *Q*_H3_), F(*Q*_H1_, *Q*_H2_) and F(*Q*_all_) for 12-10-6 potential. (G) The heat capacity curves from the ProfasiGo models with the two potentials. (H) The free energy profiles of F(*Q*_tot_) at corresponding *T_f_*. The results are from multicanonical MC simulations with the ProfasiGo model with *ϵ*_Go_=0.3. For ProfasiGo model with 12-10-6 potential we found
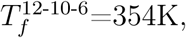 while for ProfasiGo model with 12-10 potential, 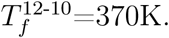

**Figure S5:**
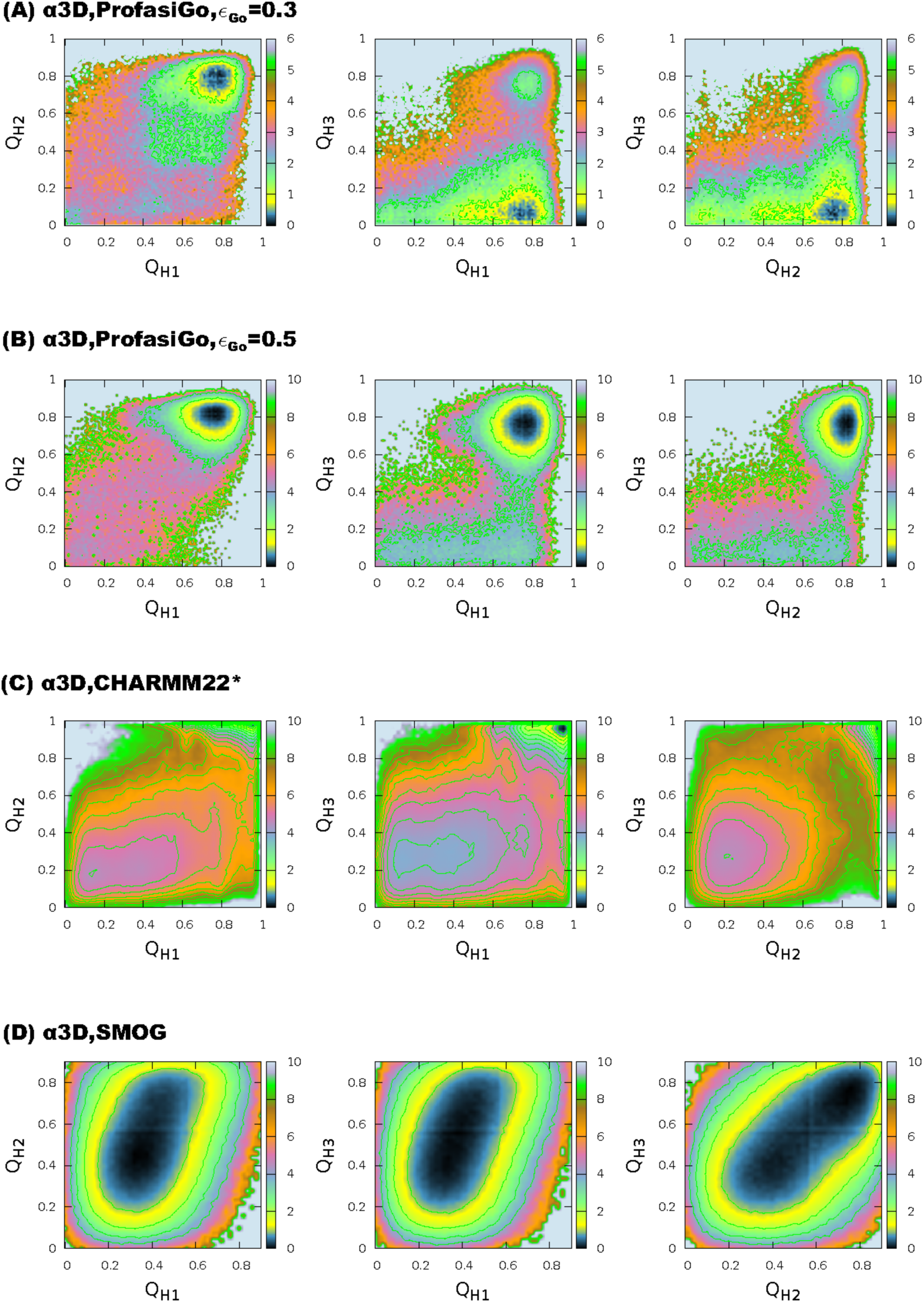
Comparison of the free energy landscapes of *α*3D obtained from the pure Gō model, the hybrid ProfasiGo model and an explicit solvent force field. (A) Free energy surfaces of *α*3D as a function of *Q*_H1_, *Q*_H2_ and *Q*_H3_ obtained from the ProfasiGo model with *ϵ*_GO_=0.3 at *T_f_*=339K. (B) Free energy surfaces of *α*3D obtained from the ProfasiGo model with *ϵ*_GO_=0.5 at *T_f_*=373K. (C)The same free energy surfaces of *α*3D obtained from a previously published 707*μ*s long MD simulation with CHARMM22^∗^ at T=370K. (D) The same free energy surfaces of *α*3D obtained from MD simulations with a pure Gō model (SMOG) at *T_f_*=124K.

**Figure S6:**
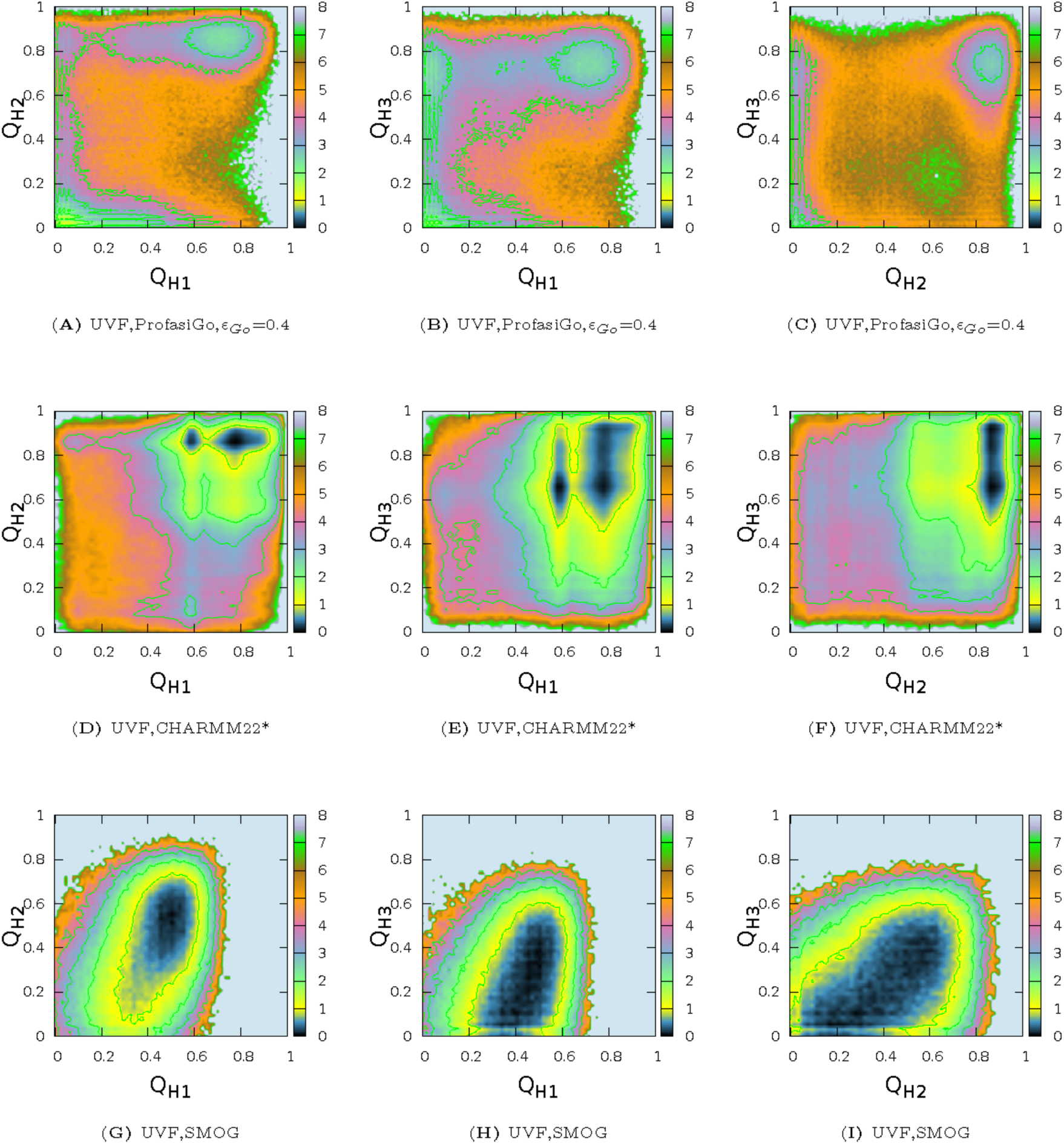
Comparison of the free energy landscapes of UVF obtained from the pure Gō model, the hybrid ProfasiGo model and an explicit solvent force field. (A–C) Free energy surfaces obtained using the ProfasiGo model with *ϵ*_G_O__=0.4 at *T_f_*. (D–F) Free energy surfaces obtained using all-atom MD with the CHARMM22^∗^ force field at 360 K, somewhat below the melting temperature in this force field (390 K). (G–I) Free energy surfaces obtained using the the pure Gō model (SMOG) at 124 K.

**Figure S7:**
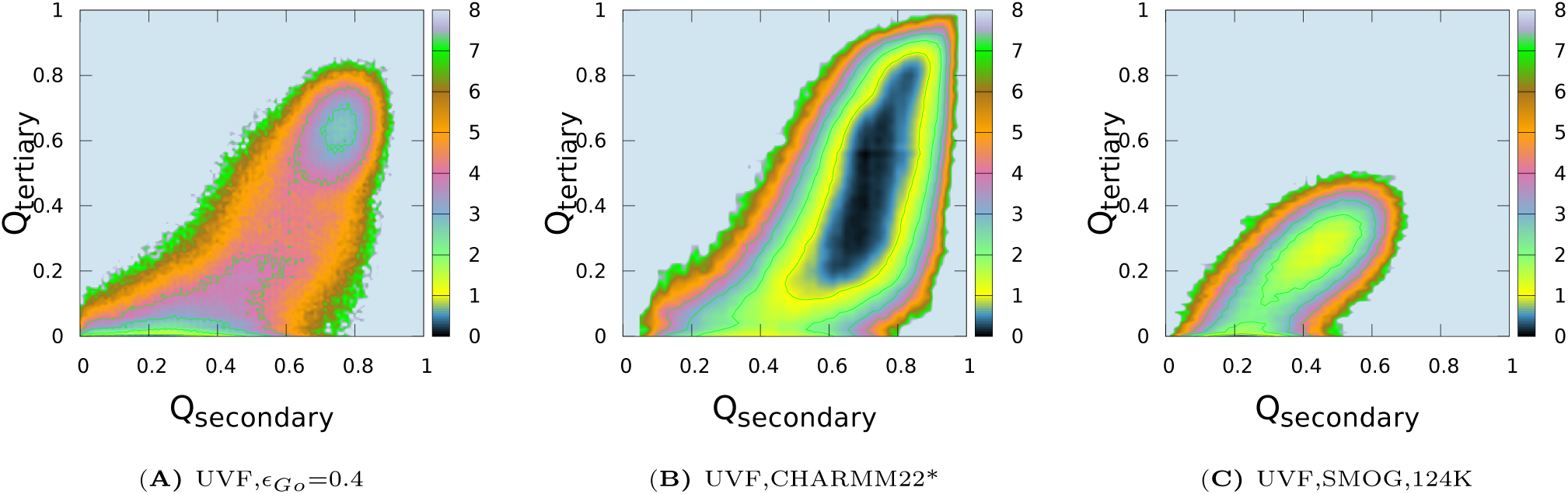
Comparison of the free energy landscapes of UVF obtained from the pure Gō model, the hybrid ProfasiGo model and an explicit solvent force field. The figure shows the free energy surfaces of UVF as a function of *Q*_Secondary_ (the fraction of native contacts within the three helices) and *Q*_tertiary_ (the fraction of native contacts between the three helices) obtained from (A) the ProfasiGo model with *ϵ*_Go_ = 0.4, (B) CHARMM22^∗^ with TIP3P water, and (C) the pure Gō model (SMOG).

